# KH1 Domain of Poly C Binding Protein 1 (PCBP1) Interacts with Utrophin-A mRNA and Modulates Utrophin-A Expression in Mouse Myoblasts

**DOI:** 10.64898/2026.10.31.763282

**Authors:** Doel Pal, Arka Bagchi, Subhanwita Das, Gargi Ghosh, Soumyak Pal, Shreya Saha, Chhanda Sarkar, Utpal Basu

## Abstract

Utrophin upregulation is an attractive strategy to ameliorate Duchenne Muscular Dystrophy (DMD). Here, we investigated whether the murine poly(C)-binding protein 1 (PCBP1) directly interacts with the utrophin-A mRNA, and how the interaction shapes the fate of utrophin-A in muscle cells. As no experimentally validated structure of the murine PCBP1 is available, AlphaFold2 was utilized to predict the structure of the full-length protein and the KH1-KH2 domain of the protein. The more stable and reliable model of the KH1-KH2 domain of PCBP1 was then docked with the 5′UTR of the utrophin-A mRNA, and the top-ranked docked complex was utilized for MD simulation to identify the most stable hydrogen bond-forming amino acid residues from PCBP1. ITC showed a thermodynamically favorable spontaneous interaction between the full-length PCBP1 and the 5′UTR of utrophin-A mRNA, but the deletion mutant of PCBP1, lacking the predicted contact region, failed to exhibit any interaction. This observation was also confirmed by an RNA-immunoprecipitation assay in C2C12 cells. However, the mutant PCBP1 was unable to alter the expression of the utrophin-A in the C2C12 cells. In cells, PCBP1 overexpression lowered utrophin-A expression, while shRNA knockdown produced reciprocal increases. A 5′UTR-luciferase reporter mirrored this observation. Finally, subcellular fractionation showed that PCBP1 promotes nuclear retention of utrophin-A transcripts and reduces their cytoplasmic availability, with reciprocal shifts upon PCBP1 knockdown. Considering earlier work on PCBP2, our data suggest both paralogs recognize the same 5′UTR element, and the KH1-dependent interaction represses utrophin-A by coupling nuclear retention to reduced protein output. Thus, the PCBP1-5′UTR interaction emerges as a potent target for utrophin de-repression in DMD.

## Introduction

Utrophin, encoded by the *UTRN* gene on chromosome 6, shares significant structural and functional homology with dystrophin. Structurally, utrophin contains an N-terminal actin-binding region, a central spectrin-repeat rod domain, and C-terminal protein-interaction domains, allowing it to associate with components of the dystrophin-associated protein complex and support sarcolemmal integrity. During embryonic development, utrophin is ubiquitously expressed at the sarcolemma; however, in postnatal and adult muscle, its expression becomes restricted primarily to the neuromuscular junctions and myotendinous junctions ([UTRN utrophin]; Deconinck et al. 1997a). Experimental studies have shown that overexpression of utrophin in dystrophin-deficient mice, either through transgenic manipulation or viral vector-mediated delivery, can ameliorate muscle pathology and improve muscle function (Tinsley et al. 1996; Deconinck et al. 1997b). Therefore, pharmacological or genetic strategies aimed at upregulating utrophin expression throughout the muscle fiber hold significant promise as a therapeutic avenue for Duchenne Muscular Dystrophy (DMD), which is the most common and severe form of X-linked muscular dystrophy (Rafael et al. 1998; Kennedy et al. 2018; Soblechero-Martín et al. 2021).

Despite this potential, the translation of utrophin-based therapies into clinical settings remains challenging due to our incomplete understanding of the regulatory mechanisms governing utrophin expression, especially at the post-transcriptional level. Among the various isoforms of utrophin, utrophin-A is the predominant form expressed in skeletal muscle (Weir et al. 2002). It is regulated by a complex array of transcriptional and post-transcriptional elements, including 5’ and 3’ untranslated regions (UTRs), internal ribosome entry sites, and RNA-binding proteins (RBPs) (Loro et al. 2020; Miura et al. 2005; Ghosh and Basu 2015).

Recent studies have highlighted the role of RBPs in modulating mRNA stability and translation of muscle-specific genes. One such protein, Poly(C)-binding protein 1 (PCBP1), is a multifunctional RBP known to bind C-rich elements within the 5′UTRs and 3′UTRs of target mRNAs, thereby influencing mRNA localization, stability, and translational efficiency. PCBP1 has been implicated in diverse biological processes, including iron metabolism, immune responses, and myogenesis. Moreover, previous studies have shown by molecular dynamics and energetic analyses how residue-specific contacts and RNA conformational changes can influence PCBP1 recognition (Gillet and Dumont 2022). Similarly, broader studies of PCBP1/hnRNP E1-mediated transcript regulation support the view that PCBP1 can act in a transcript-selective manner, with regulatory outcome depending on RNA context and cellular state (Tang et al. 2017). Notably, the presence of C-rich motifs within the 5′UTR of utrophin-A mRNA suggests a potential regulatory interaction with PCBP1 (Ryu et al. 2017; Makeyev and Liebhaber 2002; Espinoza-Lewis et al. 2017).

Given the therapeutic relevance of utrophin-A in DMD and the previously established role of PCBP2 in post-transcriptional regulation, it is important to investigate whether PCBP1, a closely related paralog of PCBP2, can interact with the 5′UTR of utrophin-A mRNA. Addressing this question is important as both PCBP1 and PCBP2 possess the KH domains but diverge functionally on target mRNAs (Makeyev and Liebhaber 2002). This line of inquiry may reveal novel regulatory pathways that could be exploited to enhance utrophin expression in DMD patients.

In this study, therefore, we investigated the interaction between PCBP1 and the utrophin-A mRNA. We also aim to identify the specific region of the PCBP1 that is responsible for the interaction with the 5′UTR of utrophin-A mRNA and whether modification of the specific region of the PCBP1 influences utrophin-A expression in cells. A comprehensive understanding of this regulatory axis may pave the way for RNA-targeted strategies for DMD therapy.

## Results

### Modelling of Full-length PCBP1 and Its KH1 Domain

Since there is no experimentally determined full-length murine PCBP1 structure available in public repositories ([UniProt]), the complete *Mus musculus* PCBP1 sequence was modelled with AlphaFold2-Colabfold to enable *in-silico* interaction studies. The full-length prediction yielded a bipartite confidence profile. Amino acid residues ∼1–155 and ∼290–345 exhibited high per-residue confidence (predominantly pIDDT ≥80–90), interrupted by pronounced low-confidence segments centered in the middle of the polypeptide. The PAE heatmaps for the top five ranked models (Figure 1A) showed consistently low intra-segment errors within the N-terminal (∼1–155) and C-terminal (∼290–345) regions, but high inter-segment errors between them, implying that while local domain topologies are well defined, their relative orientation is uncertain. Specifically, pIDDT values dipped to 50–70 around ∼70–80 amino acid residues and ∼150–165 amino acid residues, which were consistent with flexible loops. Again, the pIDDT fell below ∼50 from ∼170 to ∼280 amino acid residues, indicating an extended intrinsically disordered or poorly constrained linker (Figure 1B). This pattern coincided with the multiple-sequence-alignment coverage, which was dense over the termini and sparse across ∼170–280, reinforcing the interpretation that the central tract lacks strong evolutionary constraints and may be conformationally heterogeneous (Figure 1C). The PDB structure in Figure 1D shows the residue-specific pIDDT score of the best model (rank_1) of the full-length PCBP1 protein.

**Figure 1:**
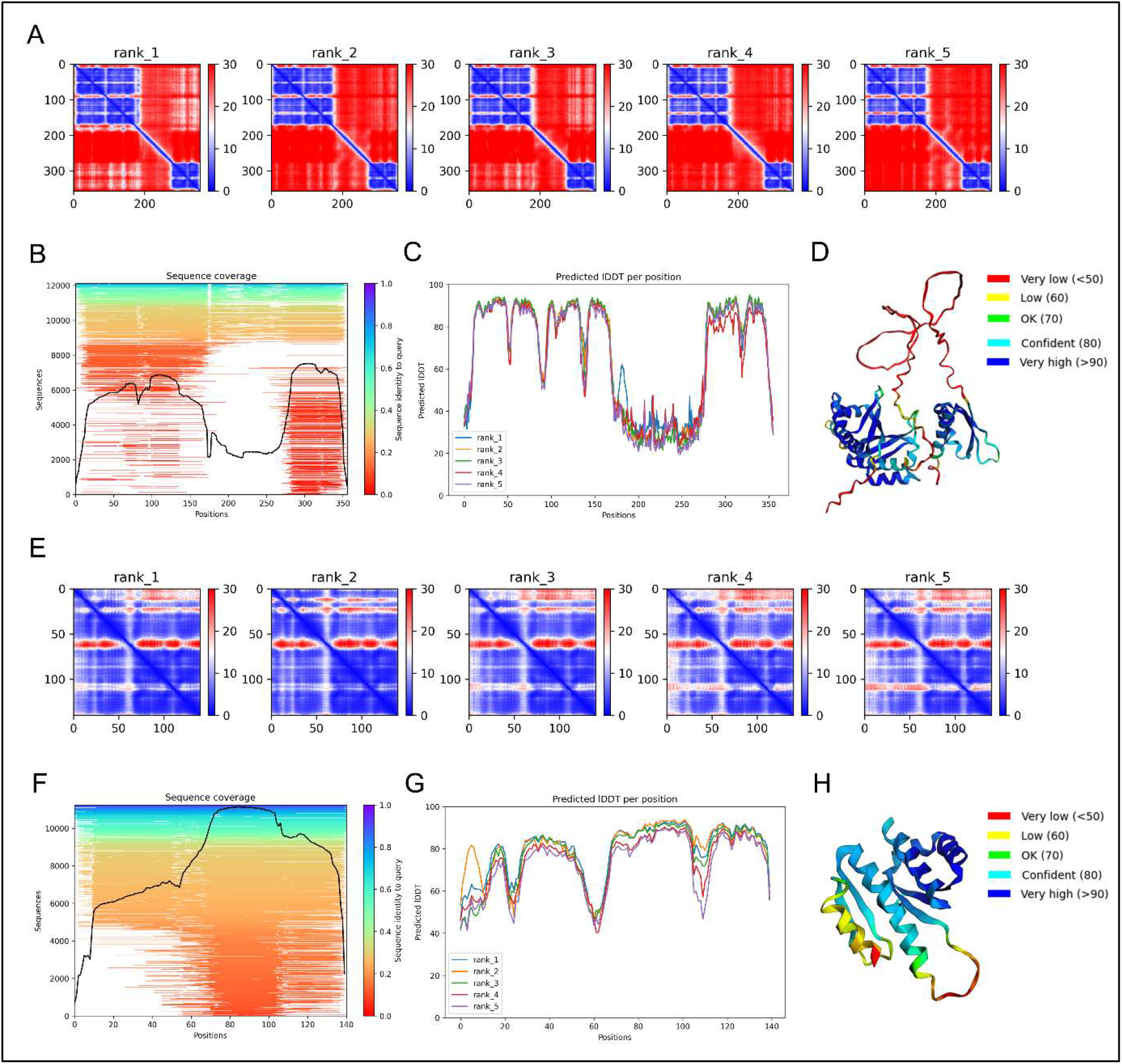
AlphaFold2 models and confidence metrics for full-length PCBP1 and its KH1 domain. (A–D) Full-length PCBP1. (A) Predicted Aligned Error (PAE) heatmaps for the top five ranked AlphaFold2 (ColabFold) models; blue indicates low pairwise alignment error (high confidence in relative placement), red indicates high error. Two well-defined blocks are evident at the N- and C-termini with high inter-block PAE, consistent with uncertain relative orientation of domains. (B) Sequence-coverage plot showing multiple-sequence-alignment (MSA) depth along the sequence. Coverage is high over the termini and reduced across the central tract. (C) Per-residue confidence (pIDDT) plot; residues ∼1–155 and ∼290–345 score predominantly in the confident/very-high range (pIDDT ⪎80–90), whereas an extended region ∼170–280 falls below ∼50, consistent with intrinsic disorder or a flexible linker. (D) 3D model of the full-length protein colored by pIDDT (dark blue: pIDDT >90; Cyan: 80; green: 70; yellow: 60; orange/red: <50), illustrating high-confidence terminal folds separated by a low-confidence central segment. (E–H) KH1-KH2 domain of PCBP1. (E) PAE heatmaps for the top five ranked KH1-KH2 predictions showing uniformly low intra-domain PAE. (F) MSA sequence-coverage for the KH1-KH2 construct. (G) pIDDT profile demonstrating globally high confidence across the KH1-KH2 ββααβ core (minor decreases only at extreme ends). (H) KH1-KH2 3D model colored by pIDDT, highlighting a compact, well-defined fold suitable for docking.

Molecular docking of this full-length PCBP1 with the 5′UTR of the utrophin-A mRNA was performed to assess the possible interaction between the protein and the mRNA. The top-ranked docked model, obtained from the HDOCK server, showed a strongly favorable docking score of -317.45, but with an unrealistically high backbone root-mean square deviation (RMSD) of 145.17Å (Figure S1). This behavior was consistent with the confidence profile of the flexible, low-pIDDT segments and the uncertain inter-domain geometry, obtained from AlphaFold2. Importantly, the mapped contacts between the protein and the mRNA in the top-ranked docked pose clustered almost entirely within amino acid residues 11-75. These residues specifically reside under the KH1 domain of the mouse PCBP1 that also displays high local confidence in the full-length prediction. These observations indicate that the apparent scoring advantage arises from a compact, well-folded KH1 domain, whereas the poorly constrained remainder of the chain contributes noise to pose comparison.

Therefore, to obtain a construct with uniformly high local confidence and improve the docking convergence and interpretability, a portion of the PCBP1 (11 to 169 amino acid residues) was modelled separately (Figure 1E–H), which contained the KH1 and KH2 domains. In contrast to the full-length run, its PAE matrix was uniformly low within the domain, indicating a well-defined, internally consistent fold (Figure 1E), and the KH1-KH2 prediction showed globally high pIDDT (largely ≥85 across the ββααβ core) with only minor terminal decreases at the extreme ends (Figure 1F and G). On the basis of these confidence metrics and the reduced inter-domain uncertainty, the KH1-KH2 model was selected for subsequent protein-mRNA docking and molecular-dynamics simulations, whereas the low-confidence central region of the full-length model was not used to constrain the interface.

### KH1 Domain of PCBP1 is Essential for Interaction with Utrophin-A mRNA

To determine the interface of interaction between the KH1-KH2 domains of murine PCBP1 protein and the 5′UTR of the utrophin-A mRNA, molecular docking studies were carried out. The top-ranked docking pose of the protein-mRNA complex obtained from HDOCK was associated with a docking score of –357.26 with a root-mean square deviation (RMSD) of 46.31 Å. According to the scoring function, this highly negative value indicates a strong and favorable interaction between the KH1-KH2 domains of the PCBP1 and the target 5′UTR of utrophin-A mRNA. While the docking score is dimensionless and not a direct measure of binding free energy (ΔG), it provides a relative estimate of interaction strength, with lower values corresponding to more stable complexes (Figure 2A). To further ascertain the stability and integrity of the docked complex, molecular dynamics simulations of the docked complex were carried out for a total of 50 nanoseconds (ns). The analysis of the trajectories during the course of the simulation revealed that there were an average of ∼35 stable hydrogen bonds formed between the KH1-KH2 domains of PCBP1 and the 5′UTR of utrophin-A mRNA throughout the simulation period (Figure 2B). Comparison of the initial and the final configuration of the complex showed that PCBP1 stayed in the same binding pocket throughout the simulation period. Analyzing the RMSD of the protein backbone revealed that the bare PCBP1 protein stabilized in water at an RMSD of 0.25 nm from the onset of the simulation; similarly, the PCBP1 also stabilized within the complex at an RMSD of 0.6 nm after 10 ns (Figure 2C). The bare RNA molecule, on the other hand, showed significantly better stability at an average RMSD of 2.5 nm compared to the RNA molecule in complex with PCBP1 protein (Figure 2D). The analysis of the amino acid or nucleic acid residue-specific root-mean square fluctuations (RMSF) clearly demonstrated that both PCBP1 and the 5′UTR of utrophin-A mRNA showed significant structural alterations while in complex, compared to the unbound states (Figure 2E and F). The analysis of the radius of gyration of the protein, RNA, and the complex also ascertained the compactness of the structures in both unbound as well as bound conditions, indicating structural stability and integrity (Figure S2A and S2B). The analysis of the hydrogen bond formation revealed that the most frequently and stably interacting amino acid residues from the KH1 domain of PCBP1 protein, throughout the course of the simulation, were 23^rd^ Lysine (K), 27^th^ and 35^th^ Serine (S), 37^th^ Lysine, 39^th^ Isoleucine (I), 40^th^ Arginine (R), and 41^st^ Glutamic acid (E). Interestingly, all these amino acids are situated within the KH1 domain of the PCBP1 protein, which spans from the 13^th^ Threonine residue to the 75^th^ Isoleucine residue (Figure 2G).

**Figure 2:**
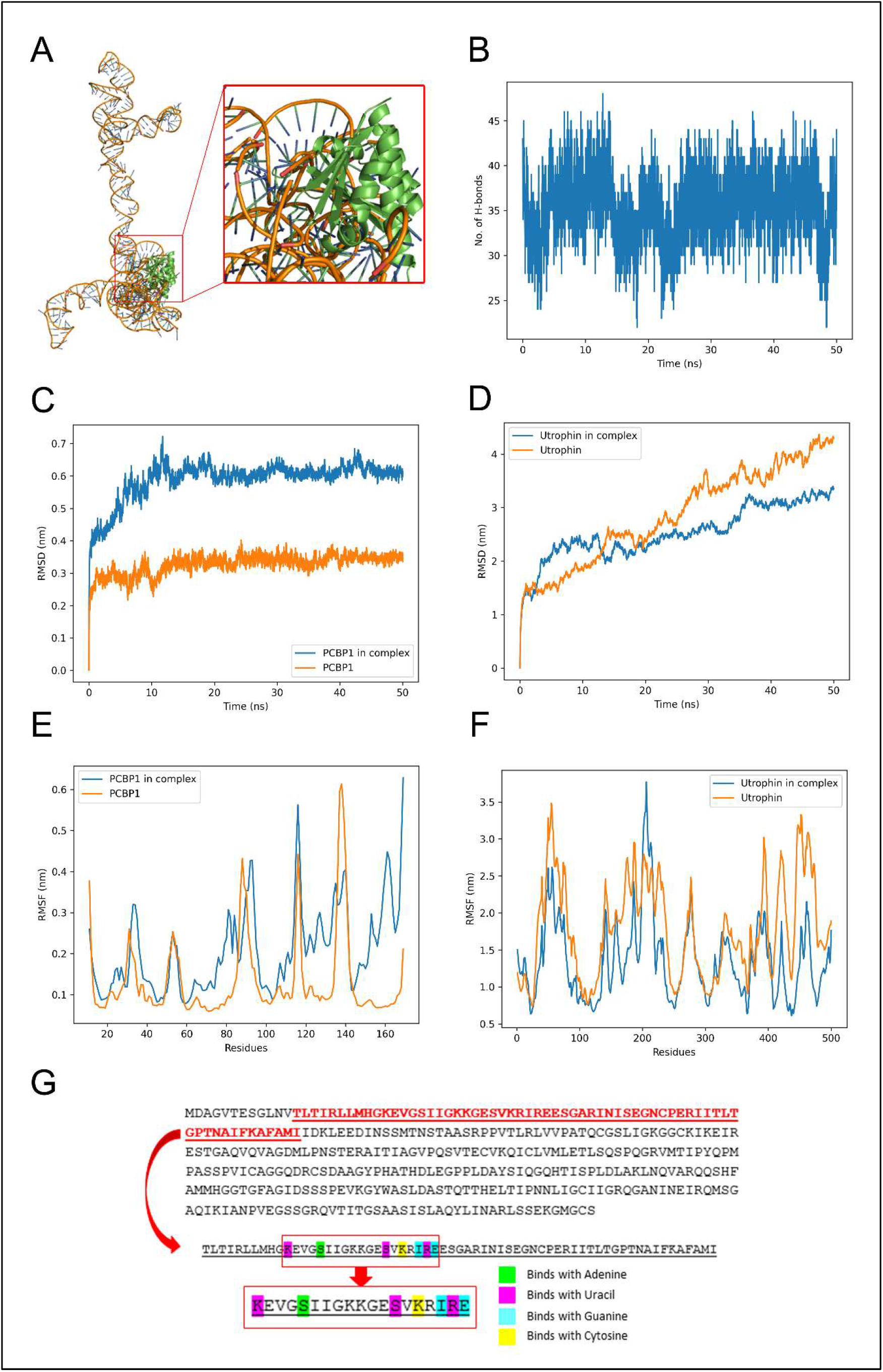
Identification of interaction sites between PCBP1 and utrophin-A 5′UTR. (A) The most favorable docked pose between the KH1-KH2 domain of PCBP1 protein and the utrophin-A 5′UTR, which showed the highest binding score. (B) Graphical representation showing the number of hydrogen bonds formed between the KH1-KH2 PCBP1 and the utrophin-A 5′UTR during the 50 ns simulation period. The line graph showing the root-mean square deviation (nm) versus time (ns) of the free PCBP1 and PCBP1 in complex with utrophin-A 5′UTR (panel C), and of the free utrophin-A 5′UTR and utrophin in complex with PCBP1 (panel D). (E) and (F) shows the residue-specific root-mean-square fluctuations of the PCBP1 protein and utrophin 5′UTR, respectively, in their free form and in complex. (G) Amino acid sequence of the full-length mouse PCBP1 obtained from NCBI, where the red-highlighted portion represents the KH1 domain. The amino acid sequence shown in the red box panel is deleted in the delPCBP1 mutant.

### Biophysical Characterization of PCBP1–Utrophin-A mRNA Interaction

To confirm a direct physical interaction between PCBP1 protein and the 5′UTR of utrophin-A mRNA, Isothermal Titration Calorimetry (ITC) was performed using purified recombinant PCBP1 (Figure S3A) and *in vitro* transcribed 5′UTR of utrophin-A mRNA. The thermodynamic profile revealed a binding isotherm characteristic of a specific molecular interaction. The upper panel of Figure 3A displays the heat pulses (DP in μcal/s) generated upon sequential injections of the RNA into the protein solution over a 50-minute run, while the lower panel shows the integrated binding heats (ΔH in kcal/mol) plotted against the molar ratio of RNA to protein (Figure 3A).

**Figure 3:**
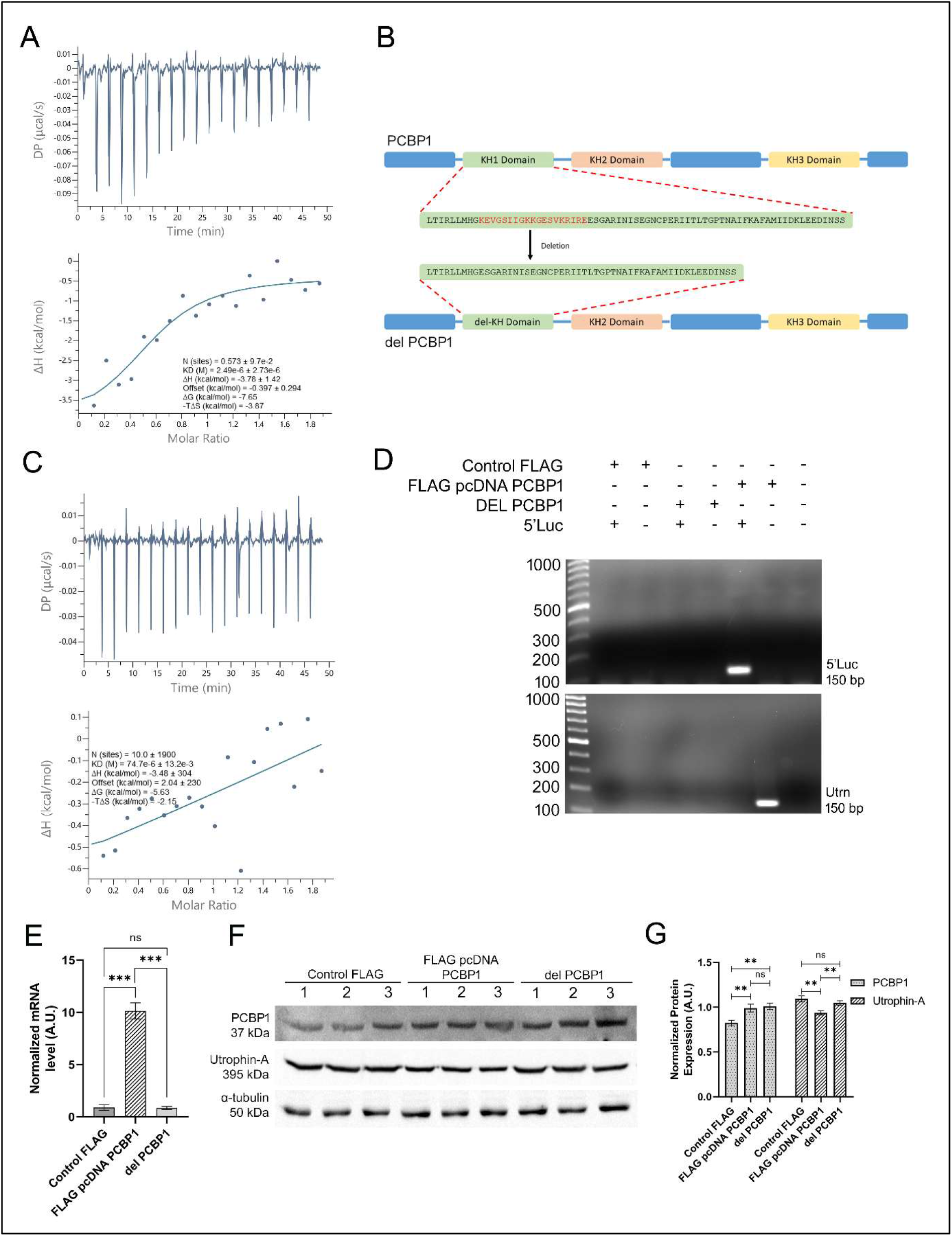
KH1 domain of PCBP1 directly interacts with utrophin-A 5′UTR. (A) Upper panel: Raw ITC thermogram showing differential power (μcal/s) over time during titration of utrophin-A 5′UTR RNA into recombinant PCBP1. Lower panel: Binding isotherm (ΔH vs. molar ratio) fitted to a one-site binding model. The dissociation constant (K_D_) was ∼2.49 μM, with a binding stoichiometry (N) of 0.573. Thermodynamic parameters indicate exothermic and enthalpy-driven binding: ΔH = –3.78 kcal/mol, ΔG = –7.65 kcal/mol, TΔS = –3.87 kcal/mol. (B) Illustration of the full-length PCBP1 containing the KH1 domain and the KH1 domain-deleted PCBP1 (delPCBP1). The deleted region (KEVGSIIGKKGESVKRIRE) within the KH1 domain is shown in red. (C) Representative ITC thermogram (upper panel) and binding isotherm (lower panel) reveal severely impaired and nonspecific binding of delPCBP1 to the utrophin 5′UTR. Key parameters (N, KD, ΔH, ΔG, TΔS) indicate a drastic loss of affinity compared to full-length PCBP1. (D) PCR analysis of FLAG immunoprecipitated RNA from cells co-transfected with FLAG-HA vector variants (Control FLAG, FLAG pcDNA PCBP1, and delPCBP1) and 5′Luc. 5′Luc-specific bands are seen only with full-length PCBP1 (FLAG pcDNA PCBP1). Similarly, endogenous utrophin-A RNA is enriched exclusively with full-length PCBP1, not with delPCBP1. (E) Bar diagram representation showing normalized mRNA levels of pcbp1 in C2C12 cells following transfection with Control FLAG, FLAG pcDNA PCBP1, and the pcDNA delPCBP1 (deletion construct). (F) Western blot analysis showing expression of PCBP1, utrophin-A, and α-tubulin following transfection of Control FLAG (empty backcone construct), FLAG pcDNA PCBP1 (full-length PCBP1), and delPCBP1 in C2C12 cells. Lanes 1, 2, and 3 represent three independent biological replicates. (G) Corresponding densitometric quantification indicates delPCBP1 was robustly expressed but did not significantly change in utrophin-A expression under this condition. These data are consistent with a requirement for the deleted KH1 region in PCBP1-dependent utrophin-A regulation. ‘ns’ means *p* > 0.05 and ** means *p* < 0.001.

Fitting the data to a one-site binding model yielded a dissociation constant (K_D_) of ∼2.49 μM, indicating moderate-affinity binding between PCBP1 and the utrophin-A 5′UTR. The stoichiometry value (N) was calculated as ∼0.573, suggesting that one PCBP1 molecule likely interacts with a partial or structured region of the RNA molecule, a feature commonly observed in RNA-binding proteins that recognize secondary structures or multi-part motifs. The binding interaction was thermodynamically favorable, with an enthalpy change (ΔH) of –3.78 kcal/mol and a free energy change (ΔG) of –7.65 kcal/mol, indicating spontaneous binding under physiological conditions. The negative entropy contribution (TΔS = –3.87 kcal/mol) suggests a reduction in disorder upon complex formation, consistent with the formation of specific RNA-protein contacts and potential stabilization of RNA structure upon binding. Together, these data confirm that PCBP1 directly binds the utrophin-A 5′UTR with micromolar affinity.

### Deletion in KH1 Domain Abrogates RNA-Binding Ability of PCBP1

To further validate the role of the KH1 domain of PCBP1 in mediating the interaction of PCBP1 with utrophin-A mRNA, a deletion construct lacking a critical region of the KH1 domain was generated. A partial deletion was made within the conserved KH1 domain. Based on the predicted interactions obtained from the molecular docking and MD simulation studies, the predicted region essential for nucleic acid binding was removed (amino acid sequence: KEVGSIIGKKGESVKRIRE, Figure 3B). The truncated variant, hereafter referred to as delPCBP1 (Figure S4B), was cloned into the FLAG-HA-tagged pcDNA3.1(+) vector (Figure S4A). The FLAG tags were utilized for immunoprecipitation.

ITC and RNA-IP were performed using the delPCBP1 mutant. For ITC, the wild type PCBP1 (Figure S3A) and the same delPCBP1 mutant (Figure S3B) were cloned into pET 28a(+) and purified. In contrast to the full-length PCBP1, which showed clear binding with a dissociation constant (K_D_) of ∼2.49 µM, ITC data for the mutant PCBP1 exhibited markedly weakened and imprecise binding characteristics (Figure 3C, upper panel). The thermodynamic parameters were as follows: N (binding sites) = 10 ± 1900, K_D_ = 74.7 µM ± 13.2 mM, ΔH = –3.48 kcal/mol ± 304, ΔG = –5.63 kcal/mol, and TΔS = –2.15 kcal/mol (Figure 3C, lower panel). The extraordinarily large error margins in stoichiometry and enthalpy, alongside a significantly weaker K_D_, underscore a loss of specific interaction between delPCBP1 and the utrophin 5′UTR. This supports the hypothesis that the KH1 domain is critical for RNA recognition and stable binding by PCBP1.

Next, the C2C12 cells, which express utrophin-A, were transiently transfected with the empty FLAG-HA vector (Control FLAG) or full-length PCBP1 (FLAG pcDNA PCBP1) or delPCBP1. To further confirm the RNA-binding efficiency of the KH1-deleted mutant, RNA-IP assays were conducted using anti-PCBP1 antibody in C2C12 cells co-transfected with FLAG-HA vector variants (Control FLAG, FLAG pcDNA PCBP1, or delPCBP1) along with 5′Luc (a luciferase construct containing the utrophin-A 5′UTR upstream of the firefly luciferase gene). RT-PCR amplification of the immunoprecipitated RNA revealed that 5′Luc RNA was only detected in the FLAG pcDNA PCBP1 + 5′Luc co-transfected group, but not in delPCBP1 or vector controls. Similarly, endogenous utrophin-A RNA was only pulled down by full-length PCBP1, with no detectable enrichment in delPCBP1 samples (Figure 3D).

Furthermore, the expression levels of the utrophin-A were also validated in the C2C12 cells, following transfection with the empty FLAG-HA vector (Control FLAG) or full-length PCBP1 (FLAG pcDNA PCBP1) or delPCBP1. The qPCR analysis showed that the *pcbp1* mRNA levels increased significantly in FLAG pcDNA PCBP1-transfected cells compared to the Control FLAG-transfected cells. However, no significant change was observed in the mRNA levels of PCBP1 in delPCBP1 mutant cells (Figure 3E). The qPCR primer pair (FP: 5′-CGGAAAGGAAGTAGGCAGCAT-3′, RP: 5′-GCAGTTCCCCTCCGAGATG-3′) used in this assay maps to the region deleted in delPCBP1 and therefore detects full-length PCBP1, or endogenous PCBP1, but does not quantify the delPCBP1 transcript. Thus, delPCBP1 expression was assessed at the protein level by immunoblotting, where significantly increased expression of the truncated protein was observed. Interestingly, no significant change in utrophin-A protein expression was observed upon delPCBP1 overexpression (Figure 3F and G).

### Overexpression of PCBP1 Alters Utrophin-A Expression

Since the mutant PCBP1 was unable to exert any effect on the protein-level expression of utrophin-A, it was essential to observe the fate of utrophin-A upon overexpression and knockdown of the full-length PCBP1. The full-length coding sequence of *PCBP1* was cloned into the mammalian expression vector pcDNA3.1(+), using *EcoRI* and *HindIII* restriction sites (Figure S5A). The integrity of the construct (pcDNA-PCBP1) was confirmed by restriction digestion and Sanger sequencing (Figure S5B). The recombinant plasmid was transfected into C2C12 myoblasts to achieve ectopic overexpression of PCBP1. qPCR data showed a robust increase in *pcbp1* transcript levels compared to empty vector controls (Figure 4A), confirming successful overexpression of PCBP1. Western blot analysis also showed a ∼6-fold increase in the protein-level expression of PCBP1. Interestingly, the protein-level expression of utrophin-A was observed to be decreased by ∼2-fold compared to the empty vector-transfected cells (Figure 4B and C).

**Figure 4:**
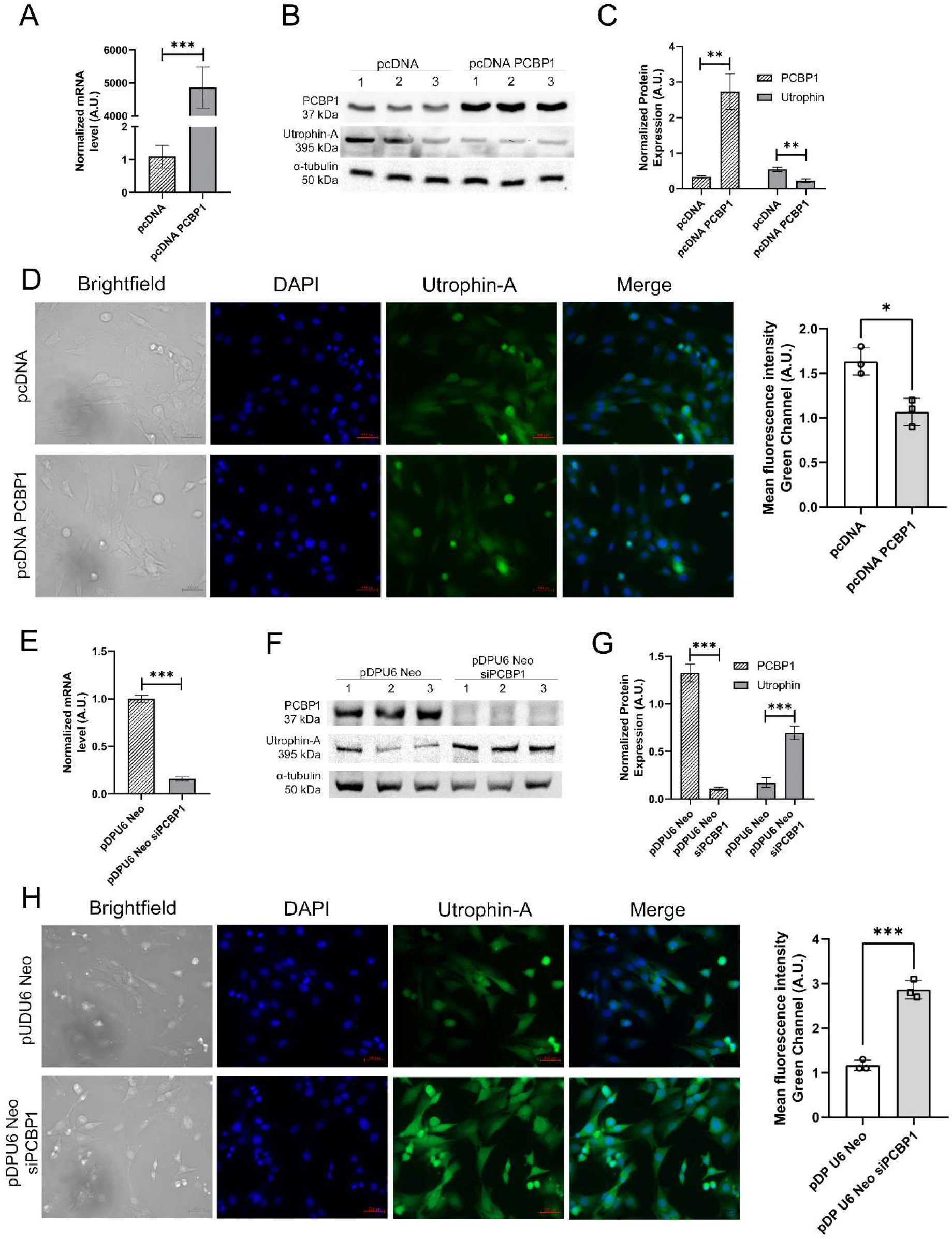
PCBP1 overexpression and silencing modulate utrophin-A expression. Assessment of the mRNA levels of pcbp1 and utrophin-A, shown in (A), revealed a significant increase in the pcbp1 mRNA levels in the C2C12 mouse myoblast cell line upon PCBP1 overexpression with the pcDNA-PCBP1 construct. The western blot analysis shown in (B) and the corresponding densitometric analysis shown in (C) demonstrated a ∼3-fold increase in the levels of PCBP1 and a ∼2-fold decrease in the utrophin-A protein levels. Lanes 1, 2, and 3 in the (B) are three biological replicates of the same experiment. Fluorescence and phase contrast microscopic images in (D) showing the intracellular expression and localization of utrophin-A protein in empty pcDNA-transfected C2C12 cells (upper panel) and pcDNA-PCBP1-transfected C2C12 cells (lower panel). (E) The normalized mRNA levels of pcbp1 in PCBP1 shRNA-expressing stable C2C12 cells, shown in the bar plots, depict a significant decrease in the pcbp1 mRNA level upon knockdown with pDPU6 Neo siPCBP1. (F) Western blot images showing the change in the protein level expressions of PCBP1 and utrophin-A of the three biological replicates in lanes 1, 2, and 3 after PCBP1 knockdown. (G) Bar plots showing densitometric analysis corresponding to the western blot of PCBP1 and utrophin-A. (H) Fluorescence and phase contrast microscopic images showing the intracellular expression and localization of utrophin-A protein in empty pDPU6-Neo-transfected C2C12 cells (upper panel) and PCBP1 shRNA-expressing stable C2C12 cells (lower panel). Data shown are from at least three independent experiments; error bars represent mean ± SEM. ** means *p* < 0.001 and *** means *p* < 0.0001.

Similar protein-level down-regulation of utrophin-A was also ascertained by immunofluorescence in C2C12 cells (Figure 4D).

These results indicated that PCBP1 plays an important role in the modulation of the protein-level expression of the utrophin-A.

### shRNA-mediated silencing of PCBP1 modulates Utrophin-A expression in C2C12 cells

Given the prior evidence that the PCBP1 protein not only interacts with utrophin-A mRNA *in vitro* but also modulates its expression in cells, knockdown of the PCBP1 was performed in the C2C12 cells to validate the regulatory effect of the PCBP1 on the utrophin-A expression. The pDPU6-Neo-siPCBP1 construct (Figure S7A-D) enabled stable expression of shRNA under the U6 promoter and selection of transfected cells using G418. This generated stable C2C12 cells that constitutively express PCBP1 shRNA. The engineered cells provided the foundation for subsequent knockdown experiments to evaluate the functional effects of PCBP1 knockdown on *utrophin-A* expression.

To assess knockdown efficiency and the resulting effect on utrophin-A expression, qPCR and Western blot analyses were performed from PCBP1 shRNA-expressing stable C2C12 cells. qPCR results revealed a substantial (>90%) downregulation of PCBP1 transcripts (Figure 4E), confirming the effective knockdown of PCBP1. Consistently, Western blot analysis also showed a significant reduction in PCBP1 protein levels by >90% (Figure 4F and G). Immunofluorescence of C2C12 cells with anti-utrophin-A antibody also showed a significant increase in the intracellular protein levels, particularly in the perinuclear region (Figure 4H).

These results demonstrated that knockdown of PCBP1 led to significant upregulation of utrophin-A at the protein level, suggesting a repressive role of PCBP1 in the regulation of utrophin-A.

To further ascertain the post-transcriptional regulation of utrophin-A mRNA by PCBP1, the 5′UTR of the utrophin-A mRNA was cloned upstream of the firefly luciferase gene (5′Luc, Figure 5A), and the activity of the gene was observed under PCBP1 overexpression and knockdown conditions. The assay showed a significant downregulation in the luciferase activity in 5′Luc-transfected cells under conditions of PCBP1 overexpression (Figure 5B). Similarly, a significant increase in the luciferase activity in 5′Luc-transfected cells was evident in PCBP1 shRNA-expressing stable C2C12 cells (Figure 5C). In contrast, no significant alteration was observed in cells transfected with the reporter gene backbone construct (pGL3-Control).

**Figure 5:**
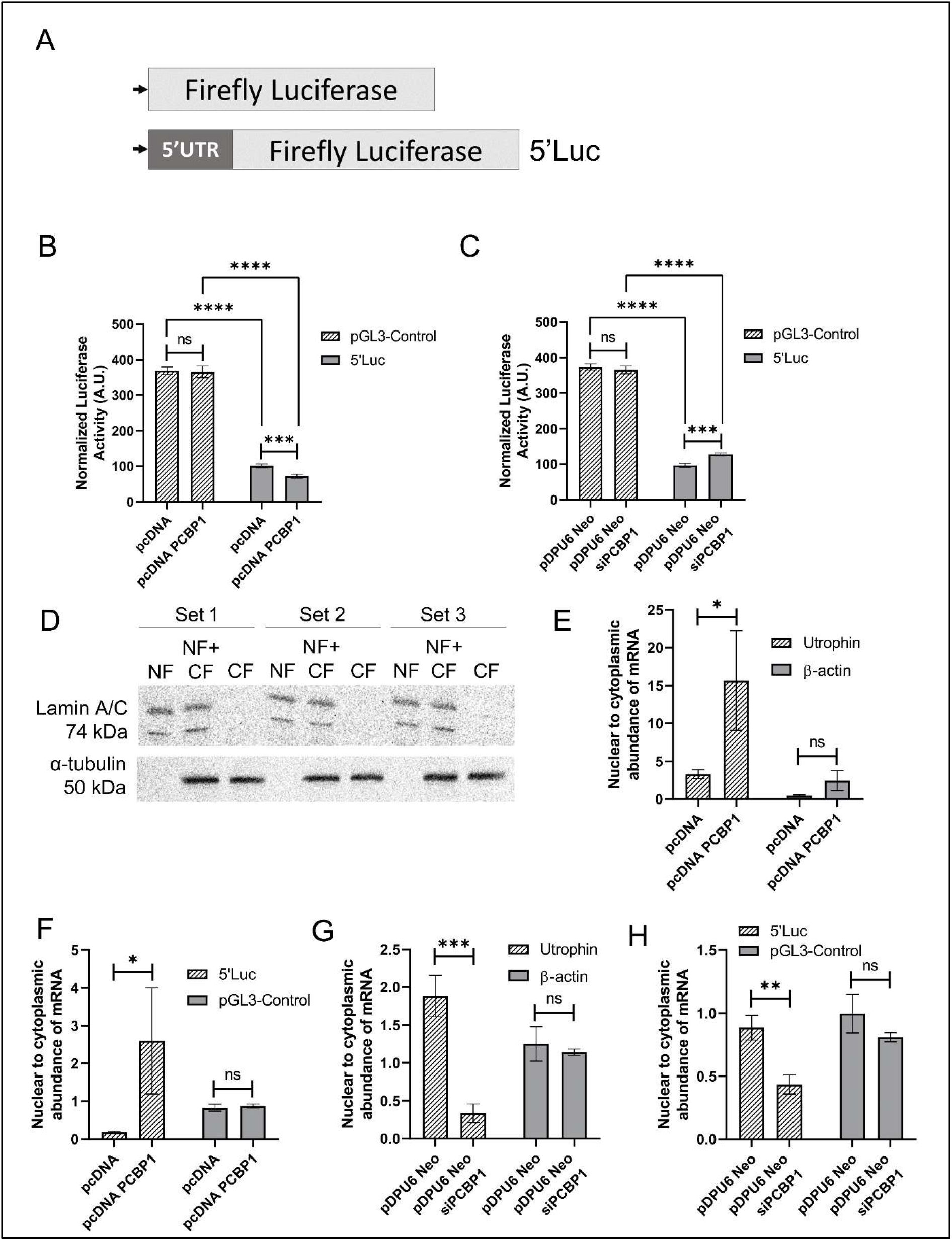
PCBP1 regulates nuclear retention of utrophin-A and 5′Luc mRNAs. (A) Schematic representation of reporter constructs pGL3-Control and 5′Luc. Bar diagram representations of the luciferase assay results in pGL3-Control or 5′Luc-transfected PCBP1-overexpressed (B) and PCBP1 shRNA-expressing stable C2C12 cells (C). (D) Western blot validation of nuclear (Lamin A/C) and cytoplasmic (α-tubulin) fractions in C2C12 cells. Sets 1, 2, and 3 represent three independent biological replicates. (E) qPCR analysis showing >4-fold increase in nuclear retention of endogenous utrophin-A mRNA upon PCBP1 overexpression; β-actin used as control. (F) 5′Luc reporter mRNA showed >8-fold nuclear retention upon PCBP1 overexpression; pGL3-Control used as control. (G) Knockdown of PCBP1 in PCBP1 shRNA-expressing C2C12 cells resulted in >8-fold decrease in nuclear retention of utrophin-A mRNA. (H) 5′Luc nuclear enrichment was reduced to <3-fold in PCBP1 shRNA-expressing cells. Data shown are from at least three independent experiments; error bars represent mean ± SEM. ‘ns’ means *p* > 0.05, * means *p* < 0.05, ** means *p* < 0.001, and *** means *p* < 0.0001.

### PCBP1 Regulates Nuclear-Cytoplasmic Distribution of Utrophin-A

We have previously reported that PCBP2 retains utrophin-A mRNA in the nucleus and therefore makes it less available for translation in the cytosol (Ghosh et al. 2021). In the current study, we, therefore, asked whether PCBP1, the closely related paralog of PCBP2, retains utrophin mRNA in the nucleus. To explore whether PCBP1 influences the subcellular localization of utrophin-A mRNA, we examined the nuclear-to-cytoplasmic distribution of endogenous utrophin-A mRNA and 5′Luc under conditions of PCBP1 overexpression and knockdown. To validate the efficacy of the nuclear-cytoplasmic fractionation protocol, western blot analysis was first performed using Lamin A/C (nuclear marker) and α-tubulin (cytoplasmic marker). Lamin A/C was exclusively detected in the nuclear fraction, while α-tubulin was restricted to the cytoplasmic fraction, confirming effective separation of compartments (Figure 5D). The nuclear and cytoplasmic abundance of the utrophin-A was analyzed by calculating 2^-ΔΔCT^, where ΔΔCT = [(C_T Nuclear (Target)_ – C_T Nuclear (GAPDH)_) – (C_T Cytosolic (Target)_ – C_T Cytosolic (GAPDH)_)], where GAPDH was used as an endogenous control.

In PCBP1-overexpressing cells, a significant increase in nuclear retention of both the 5′UTR of utrophin-A mRNA and the 5′Luc reporter was observed. qPCR revealed >4-fold enrichment of utrophin-A mRNA in the nuclear fraction relative to cytoplasmic levels, whereas no significant change in the nuclear-to-cytoplasmic abundance of β-actin mRNA was observed (Figure 5E). Similarly, 5′Luc mRNA showed a >8-fold increase in nuclear retention. Similar to that of β-actin, no significant change was observed in the nuclear-to-cytoplasmic abundance of the pGL3-Control vector mRNA (Figure 5F). These data suggest that PCBP1 overexpression promotes nuclear retention of both endogenous utrophin-A mRNA and its 5′UTR-containing reporter transcript.

Conversely, in PCBP1 shRNA-expressing stable C2C12 cells, we observed a marked reduction in nuclear retention. The nuclear-to-cytoplasmic ratio of utrophin-A mRNA dropped by more than 8-fold (Figure 5G). Similarly, 5′Luc mRNA showed more than 3-fold cytoplasmic enrichment compared to the control (Figure 5H). These findings supported the role of PCBP1 in regulating mRNA nuclear export, likely via interactions with the 5′UTR of utrophin-A.

## Discussion

Since PCBP2 has been shown to modulate utrophin-A expression and bias its nuclear- to-cytoplasmic distribution (Ghosh et al. 2021), it was essential to investigate whether the closely related paralog PCBP1 also engages utrophin-A directly, and with what consequence. Therefore, our approach began with structural modeling of the mouse PCBP1 protein, as the specific RNA-contacting surface of PCBP1 had not been resolved experimentally. AlphaFold2 predictions of full-length PCBP1 provided a confident, domain-resolved fold for KH1-KH3; however, substantial regions carried low per-residue confidence, limiting the reliability of the full-length model for complex docking. Consistently, the subsequent docking studies with the full-length PCBP1 and the 5′UTR of the utrophin-A mRNA demonstrated a significantly strong binding score, involving the majority of the amino acid residues from the KH1 domain of the PCBP1. But the unrealistically high RMSD values in the docked complex again questioned the structural integrity and reliability of the full-length PCBP1 model (Aier et al. 2016). However, it showed that the KH1 domain might be important for the interaction. Therefore, a selected portion of the PCBP1, spanning the KH1 and KH2 domains (11 to 169 amino acid residues), was remodeled using AlphaFold2, which now predicted a more reliable structural model with markedly higher residue-level confidence, providing a more suitable template for downstream analyses.

This KH1-KH2 domain of the murine PCBP1 was then utilized for the docking and MD simulation studies. Docking consistently placed a C-rich tract of the utrophin-A 5′UTR against a patch on KH1 with favorable docking energies. MD simulation for 50 ns with the most-favorable docked pose not only confirmed the stability and integrity of the complex, but also helped identify the most stable and frequent hydrogen bond-forming amino acid residues from the KH1 domain of the PCBP1.

Guided by these predictions, the protein-mRNA interaction was further validated biophysically by ITC. The ITC data supported a direct PCBP1-utrophin-A 5′UTR interaction in the micromolar range, in line with the other reported KH-RNA affinities (Yoga et al. 2012; Dagil et al. 2019; Ni et al. 2020). Simultaneously, ITC data with the delPCBP1, which lacks the specific stretch of most-interacting amino acid residues from the KH1 domain, failed to show detectable binding under similar conditions. RNA-IP with the delPCBP1-5′Luc co-transfected C2C12 cells reproduced this differential behavior, together indicating that the KH1 domain is essential for the interaction of PCBP1 with the utrophin-A mRNA. Interestingly, the deletion of the specific stretch of the KH1 domain of the PCBP1 was not able to alter the protein-level expression of the utrophin-A in the cellular environment, as validated by overexpressing the mutant PCBP1 (delPCBP1) in the C2C12 cell line.

Since it was now confirmed that the KH1 domain plays a key role in PCBP1-utrophin-A recognition, understanding how this interaction plays out in cells was of great interest. Interestingly, overexpression of the full-length PCBP1 in C2C12 myoblasts downregulated utrophin-A protein levels, whereas the PCBP1 shRNA-expressing stable C2C12 cells produced the reciprocal pattern. The luciferase reporter assays refined this view. Placing the utrophin-A 5′UTR upstream of the luciferase reporter made the reporter PCBP1-responsive, whereas the empty backbone did not affect any. This indicated that an element within the 5′UTR was responsible for the PCBP1 responsiveness.

Subsequently, the subcellular fractionation provides a link between binding and expression. Under PCBP1 overexpression conditions, both endogenous utrophin-A transcripts and the 5′Luc reporter are preferentially retained in the nucleus, while the housekeeping and backbone controls are unchanged. In contrast, in the PCBP1 shRNA-expressing stable C2C12 cells, the nuclear retention was reduced, and the cytoplasmic RNA was subsequently enriched. Because translation occurs in the cytoplasm, these shifts in partitioning offer a direct explanation for the accompanying changes in utrophin-A protein (Holt and Schuman 2013; Martin and Ephrussi 2009).

Viewed together with the earlier PCBP2 report from our laboratory, these data suggest that the two paralogs converge on the same cis-regulatory substrate, the C-rich 5′UTR of utrophin-A. In sum, the present data support an interaction mechanism, where PCBP1 binds the utrophin-A 5′UTR via KH1, biases transcripts toward the nucleus, and thereby reduces their cytoplasmic availability and protein output. This study, therefore, provides an insight into the PCBP1-mediated regulation of utrophin-A. Moreover, the alignment of human and mouse PCBP1 showed complete amino-acid identity, including the KH1 region and the deleted interaction-associated stretch (Figure S8). This supported the conservation of the protein-side component of the proposed regulatory axis. The 5′UTRs of mouse and human utrophin transcripts have also been reported to show substantial homology (Ghosh et al. 2021). However, conservation of PCBP1 does not by itself establish identical regulation in human muscle cells. Species-specific differences in utrophin-A 5′UTR architecture, RNA secondary structure, cell state, and RBP stoichiometry may influence binding and regulatory outcome. In future, it will be of great importance to define the minimal 5′UTR motif required for PCBP1 binding, confirm the contact sites on the endogenous RNA, and define the separate effects on export, decay, and validate the mechanism in human myogenic cells or DMD-derived muscle cells.

## Materials and Methods

### In silico modeling of full-length PCBP1 and the KH1 domain of PCBP1

The amino acid sequence of the mouse (*Mus musculus*) PCBP1 protein was obtained from NCBI in FASTA format (NM_011865.4). Predictions were carried out with AlphaFold2 (RRID: SCR_025454) as implemented in ColabFold (RRID: SCR_025453), using MMseqs2 to assemble multiple-sequence alignments against the default ColabFold databases. The sequence-coverage plot was exported from the run to document evolutionary information per residue. For each target, five models were generated with the default three recycles, and the models were subsequently AMBER-relaxed. All the models were ranked by per-residue confidence (pIDDT) for monomers. Pre- and post-relaxation coordinates, together with run logs, were retained.

Model quality was evaluated from the pIDDT and the Predicted Aligned Error (PAE) matrices exported for the top five ranked models, and these were interpreted in the context of the sequence-coverage plot to account for variation in MSA depth along the sequence. Contiguous segments with pIDDT ≥70 and low intra-segment PAE were treated as well-structured domains, whereas stretches with pIDDT <50 and uniformly high inter-segment PAE were interpreted as intrinsically disordered or flexibly linked. The relaxed rank_1 models were used for analysis (Kim et al. 2025; Mirdita et al. 2022).

### Molecular docking

The rank_1 model of the PCBP1 KH1 domain, generated from AlphaFold2, was used for the molecular docking studies with the 5′ untranslated region (5′UTR) of utrophin-A mRNA. The nucleotide sequence of the 5′UTR of the mouse utrophin-A mRNA was obtained from NCBI and was used to predict the secondary structure by the RNAfold web server ([RNAFold Web server]; Gruber et al. 2008). The output from the RNAfold (RRID: SCR_024427) web server (in dot-bracket format) was used as an input in the RNAComposer (RRID: SCR_026636) web server to generate the three-dimensional structure of the 5’UTR of the utrophin-A mRNA in protein data bank (PDB) format UTRN Utrophin]; Biesiada et al. 2016). Protein–mRNA docking was carried out using the HDOCK server (RRID: SCR_024799), which employs a hybrid strategy combining template-based modeling and *ab initio* free docking. The quality of docking was assessed using the server’s statistical potential scoring function (ITScore), where more negative values indicate stronger predicted binding affinity (Yan et al. 2020, 2017).

### Molecular dynamics simulation of protein-mRNA complex

Molecular dynamics (MD) simulation of the protein-mRNA docked complex was performed using GROMACS (version 2022.4, RRID: SCR_014565) (Lindahl 2008; Abraham et al. 2015) on a High-Performance Computing Cluster at the S. N. Bose Innovation Centre at the University of Kalyani. The coordinates of the protein-mRNA complex were converted into simulation-appropriate format using the AMBER force field ([The AMBER Forcefield]). The final configuration file included protein-mRNA complex, solvent simple point-charge water molecules, and charge-neutralizing ions. The stability and integrity of the protein-mRNA complex were analyzed after 50 nanoseconds (ns) by calculating the root-mean-square deviation (RMSD) and radius of gyration (Rg) of the protein and mRNA in complex, as well as individually in solvent. The structural fluctuation due to the interaction was also evaluated by calculating the root-mean square fluctuations (RMSF) of the amino acid residues of the protein. The MDTraj Python module was utilized to determine the hydrogen bond-forming atoms from the protein and the mRNA during the 50 ns simulation period (McGibbon et al. 2015). This module utilizes the Baker-Hubbard method to identify the hydrogen bonds in the trajectories of the simulation. According to the Baker-Hubbard criterion, a hydrogen bond, like d – h ··· a (donor – hydrogen ··· acceptor), is identified if the cut-off distance between the hydrogen and the acceptor atom is less than 0.25 nm (r_ha_ < 0.25 nm) and the angle between the donor, hydrogen, and the acceptor atoms in greater than 120° (θ_dha_ > 2π/3). For MD, backbone RMSD and minimal distances between the protein and RNA were monitored over time (Bagchi et al. 2025b).

### Pcbp1 cloning into pcDNA3.1(+)

The open reading frame (ORF) of *pcbp1* was PCR-amplified from cDNA using gene-specific primers (Forward: 5’-AATAGAATTCATGGACGCCGGTATGACTG-3’ and Reverse: 5’-AAAAAAGCTTCTAGCTGCACCCCATCCCC-3’) and PrimeSTAR® GXL DNA polymerase (Cat. No.: R050A, Takara Bio), and cloned at *EcoRI* and *HindIII* sites of pcDNA3.1(+) (Cat. No.: V79020, Invitrogen). The plasmid was named pcDNA-PCBP1.

### PCBP1 shRNA plasmid preparation

To achieve knockdown of PCBP1 in C2C12 cells, a series of plasmid constructs was designed and generated. The silencing element was initially derived from the pDG458 vector (Cat. No.: 100900, Addgene, RRID: Addgene_100900), which contains tandem *BbsI* restriction sites compatible with short hairpin RNA (shRNA) cloning. These *BbsI*-flanked sequences were subcloned along with the U6 promoter of pDG458 into the pBSU6 plasmid backbone, generating the construct termed pDPU6 (Fig. S8A–B). To facilitate antibiotic-based selection, a Neomycin resistance gene (NeoR) cassette was cloned upstream of the U6 promoter in pDPU6, resulting in the construct pDPU6-Neo (Fig. S8C). This allowed for effective screening of transfected cells under G418 selection (for 2-3 weeks). Finally, a validated shRNA target for PCBP1 (TRCN0000120938, GPP Web Portal, Broad Institute GPP Web Portal]); target 5′-GCCTACCAATGCCATCTTTAA-3′) was used to generate the hairpin cassette. Two complementary oligonucleotides bearing *BbsI*-compatible 5′ overhangs (forward: 5′-CACCGGCCTACCAATGCCATCTTTAAC-3′ + 5′-CCATGGTTAAAGATGGCATTGGTAGGCC-3′; reverse: 5′-AAACAAAAAGGCCTACCAATGCCATCTTTAA + 5′-CATGGTTAAAGATGGCATTGGTAGGCCTTTTT-3′) were 5′-phosphorylated with T4 polynucleotide kinase, annealed to produce a duplex encoding the U6-driven shRNA (sense-loop-antisense with a Pol III terminator), and ligated into *BbsI*-digested pDPU6-Neo downstream of the U6 promoter. Correct insertion and orientation were confirmed by Sanger sequencing across the U6-insert junctions, yielding the final construct, pDPU6-Neo-siPCBP1 (Figure S8D).

### Purification of His-tagged PCBP1 protein

The ORF of *pcbp1* was amplified from the pCDNA-pcbp1 plasmid using forward primer ATATGAATTCATGGACGCCGGTGTGACTGAAAGC and reverse primer ATATAAGCTTCTAGCTGCACCCCATCCCCTT, having *EcoRI* and *HindIII* restriction sites. These restriction sites were utilized for cloning the ORF into pET28a(+) (EMD Biosciences, Cat. No.: 69864-3) for expression of N-terminally 6xHis-tagged PCBP1. The pET-pcbp1 construct was transformed into *E. coli* BL21 (DE3) cells and induced with 1.5 mM IPTG. Cells were harvested and lysed in buffer containing 50 mM Tris-HCl (pH 8.0), 300 mM NaCl, 10 mM imidazole, lysozyme, β-mercaptoethanol, and protease inhibitors. The lysate was clarified by centrifugation. The supernatant was incubated with Ni-NTA agarose beads (Ǫiagen, Cat. No.: 30210) pre-equilibrated with lysis buffer at 4 °C. After binding, the beads were washed three times with wash buffer (Tris-HCl, NaCl, 40 mM imidazole), and protein was eluted with elution buffer (250 mM imidazole). Eluted protein was dialyzed using a Tub-O-DIALYZER^TM^ (G-Biosciences, Cat. No.: 786-613) in RNA-protein binding buffer (15 mM NaCl, 1 mM MgCl_2_, 10 mM Tris-HCl, pH 7.4) (Ghosh et al. 2021).

### Preparation of delPCBP1 constructs

The KH1-deleted PCBP1 construct (delPCBP1) was generated by PCR-based deletion cloning using the full-length pcDNA-PCBP1 plasmid as template. Outward-facing primers flanking the predicted RNA-interacting region of PCBP1 were used: forward primer 5′-GAGAGCGGCGCGCGGATC-3′ and reverse primer 5′-TCCGTGCATCAGCAGCCGAAT-3′. PCR amplification generated a linearized pcDNA3.1-PCBP1 product lacking a 57-nucleotide segment corresponding to the selected KH1-region deletion. The PCR product was purified and self-ligated using T4 DNA ligase to generate the 57-nt-deleted PCBP1 expression construct, termed pcDNA-delPCBP1. This construct was used for mammalian-cell expression and functional deletion experiments.

For recombinant protein expression and ITC experiments, the same deletion strategy was applied to the bacterial expression construct pET28a-PCBP1. Briefly, pET28a-PCBP1 was used as the template, and the same primer pair was used to amplify the 57-nt-deleted linear product. The purified PCR product was ligated with T4 DNA ligase to generate pET28a-delPCBP1, which was subsequently used for expression and purification of recombinant delPCBP1 protein.

### In vitro transcription of the 5′UTR region of utrophin-A mRNA

The 5′UTR region of utrophin-A mRNA was amplified by PCR from the 5′Luc vector, where the mouse utrophin-A 5′UTR was previously cloned. The forward primer containing a T7 promoter sequence (5′-ACTGCTAATACGACTCACTATAGGGTTGTGGAGTCGCCCTTCCC-3′) and a specific reverse primer (5′-AATTCCATGGCTTGAATGAGTTTCAGTATAATCCAAGCG-3′) were used. *In vitro* transcription was performed using the T7 polymerase (NEB, Cat. No.: M0251S) following the manufacturer’s protocol. The transcription reaction was incubated at 37°C for 3 hours, after which the synthesized RNA was treated with DNase I to remove template DNA. The RNA was purified using a phenol-chloroform extraction followed by ethanol precipitation, and its integrity was confirmed by agarose gel electrophoresis and quantification using a NanoDrop 2000 (Thermo Scientific) spectrophotometer (Ghosh et al. 2021).

### Isothermal titration calorimetry

Isothermal Titration Calorimetry (ITC) was employed to quantitatively assess the binding interaction of purified PCBP1 protein with *in vitro*-transcribed utrophin-A 5′UTR. Experiments were performed using a MicroCal PEAǪ-ITC instrument (Malvern Panalytical) at 25 °C. Prior to titration, both protein and RNA samples were extensively dialyzed against the same buffer (20 mM HEPES, pH 7.4, 150 mM NaCl, 1 mM DTT) to minimize heat effects from buffer mismatch. PCBP1 protein (20 μM) was loaded into the sample cell, and utrophin-A 5′UTR RNA (200 μM) was titrated in 2 μL aliquots at intervals of 150 seconds. The heat change accompanying each injection was recorded and corrected for dilution heat by subtracting control titrations of RNA into buffer. The resulting binding isotherms were fitted to a one-site binding model using MicroCal PEAǪ-ITC analysis software (RRID: SCR_023795) to determine the binding affinity (K_D_), enthalpy (ΔH), entropy (ΔS), free-energy change (ΔG), and stoichiometry (N) of the interaction (Ghosh et al. 2021; Feig 2009).

### RNA-Immunoprecipitation (RNA-IP)

1 × 10^7^ C2C12 cells were treated with formaldehyde and harvested in vanadyl ribonucleoside complex (400 μM) (NEB, Cat. No.: S1402S), supplemented with RIPA buffer. The cells were sonicated and added to Chromotek® DYKDDDDK Fab-Trap® Agarose (RRID: AB_2894836), supplemented with 2.5% BSA and RNaseOUT^TM^ Recombinant Ribonuclease Inhibitor (200 U/ml, Invitrogen, Cat. No.: 10777019). After washing with RIPA-HS buffer, the protein-RNA complex was disrupted in elution buffer. cDNA was synthesized using MMLV reverse transcriptase (NEW ENGLAND Biolabs, Cat. No.: M0253S) and random hexamer (Niranjanakumari et al. 2002). RT-PCR was performed using primers of utrophin and firefly luciferase described in Table 1.

**Table1:**
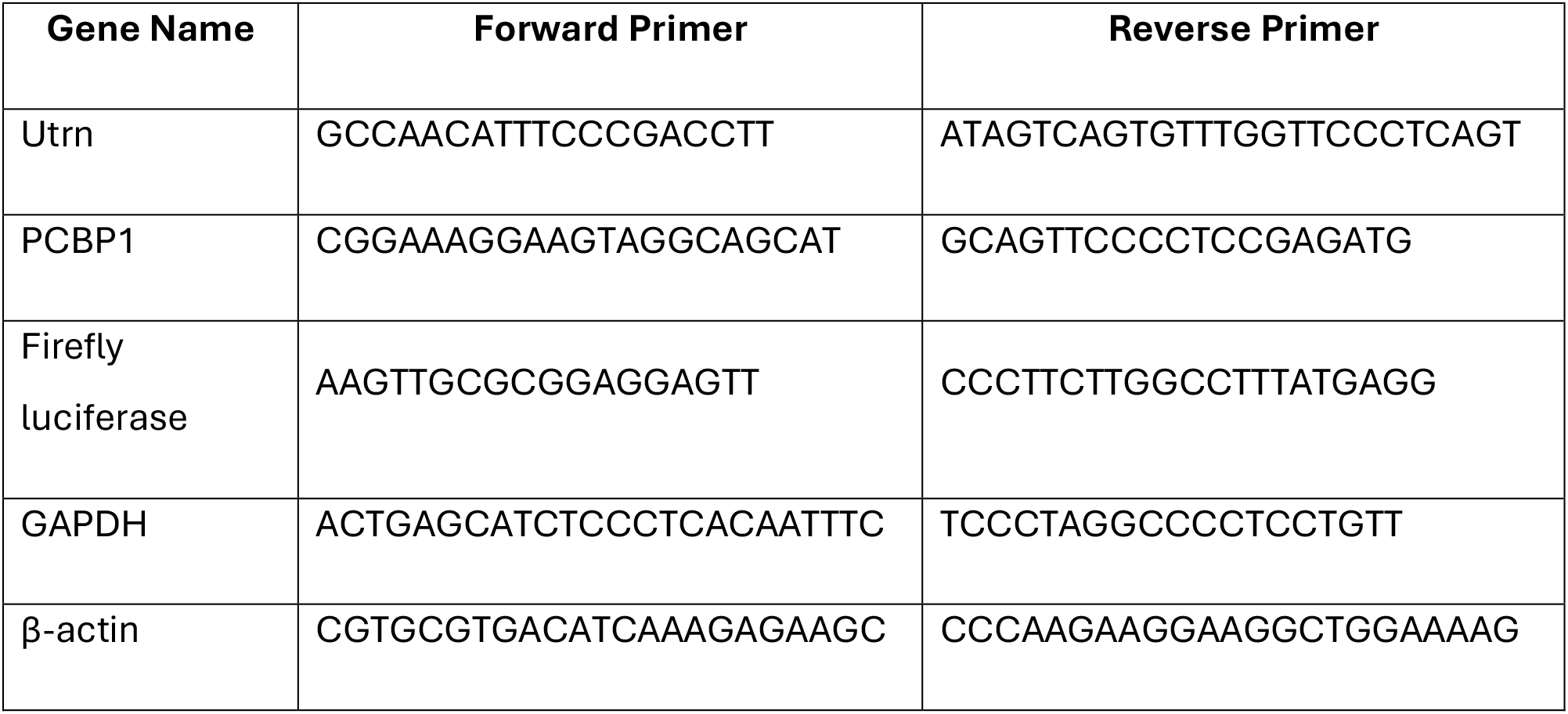
List of primers used for qPCR.

### Culture of C2C12 cell line

The mouse (*Mus musculus*) myoblast cell line C2C12 (Cat. No.: CRL-1772, RRID: CVCL_0188) was used for this study, which was obtained from ATCC, USA. The cell line was maintained in a culture medium, consisting of Dulbecco’s Modified Eagle Medium (DMEM) from Gibco^TM^ (Cat. No.: 11965092), supplemented with 5-10% of Fetal Bovine Serum (FBS) from Gibco^TM^ (Cat. No.: 11550356) and 100 U/ml of penicillin-streptomycin. The cells were kept in a T-25 sterile vented flask at 37°C. The environment was maintained with 5% CO_2_ and kept humidified (Taye et al. 2020). Before commencement of any experiment, the cells were allowed to achieve at least 70% confluence and tested for the presence of any contaminants.

### Western blot analysis

To perform a western blot, the cells were lysed with Nonidet® P-40 (NP-40) buffer (G-Biosciences, Cat. No.: DG001), and the extracted proteins were resolved using either a 12% or a 3-12% gradient polyacrylamide gel after mixing with protein loading buffer. Then the gel containing resolved proteins was soaked in transfer buffer, containing 25 mM Tris base, 190 mM glycine, and 20% methanol, and then it was transferred into a nitrocellulose (0.2μm, Bio-Rad, Cat. No.: 1620112) or an activated Immun-Blot® PVDF membrane (Bio-Rad, Cat. No.: 1620177). The membrane was then blocked with 5% BSA and incubated with primary antibody at a dilution of 1:1000 overnight at 4°C. Primary antibodies used in this study were: utrophin antibody (Cat. No.: MANCHO3(8A4), DSHB, RRID: AB_2618137), hnRNP E1 Antibody (PCBP1) (Cat. No.: 8534, Cell Signaling Technology, RRID: AB_11129258), Lamin A/C antibody (MANLAC1(4A7), DSHB, RRID: AB_2618203), and alpha-Tubulin antibody (Cat. No.: 2144, Cell Signaling Technology, RRID: AB_2210548). After primary antibody incubation, the antibody was removed, and the membrane was washed with 1X TBST buffer. Then the membrane was incubated with secondary antibody at a dilution of 1:5000 for 2 hours at room temperature. The two secondary antibodies used for this study were: Sheep Anti-Rabbit IgG Antibody – HRP conjugate (Millipore, Cat. No.: AP510P, RRID: AB_3718559) and Goat Anti-Mouse IgG – HRP conjugate (Millipore, Cat. No.: 71045-3, RRID: AB_11211441). The membrane was then subjected to immunodetection using a composition of luminol, paracoumaric acid, and hydrogen peroxide, and the image was obtained from Bio-Rad XRS+ Gel Documentation System and analyzed using Bio-Rad Image Lab software (RRID: SCR_014210) and NIH ImageJ software (RRID: 003070) (Bagchi et al. 2025a). All the replicates represent independent biological replicates.

### Immunofluorescence

Immunofluorescence detection of control C2C12 cells, PCBP1-overexpressed C2C12 cells, and PCBP1-silenced C2C12 cells was performed with primary antibodies against utrophin (Cat. No.: MANCHO3(8A4), DSHB, RRID: AB_2618137). After overnight incubation (at 4°C) with the primary antibodies, following blocking with 5% BSA, the cells were incubated with goat anti-mouse secondary antibodies conjugated with FITC (SantaCruz Biotechnology Inc., Cat. No.: sc-2010, RRID: AB_631735) for 2 hours at room temperature (Rando and Blau 1994). Following the antibody incubation, the cells were incubated with DAPI (Sigma Aldrich Cat. No.: D9542) for 15 minutes. Images were collected, after vigorous washing with 1X PBS, with a ZEISS Axio Vert A1 Microscope under identical conditions using ZEISS ZEN Microscopy Software (RRID: SCR_013672).

### q-PCR

Total RNA was isolated from C2C12 cells by the TRIzol™ (Invitrogen, Cat. No.: 15596026) method according to the manufacturer’s protocol. The isolated RNA sample was subjected to DNase I (NEB, Cat. No.: M0303S) in 1X DNase I buffer at 37°C for 30 mins. 1 µg total RNA was used to prepare cDNA. Real-Time PCR reaction was performed using iTaq ™ Universal SYBR^®^ Green Supermix (Bio-Rad, Cat. No.: 1725120), 300nM of each primer, and 0.5 µl cDNA in each well according to the manufacturer’s protocol in the Applied Biosystems StepOne™ machine (RRID: SCR_023455) (Ghosh et al. 2021). The forward and reverse primers for the q-PCR analyses are listed below (Table 1):

Ct values were called with StepOne™ software (RRID: SCR_014281) using automatic baseline/threshold settings applied uniformly across a plate. Relative transcript abundance was calculated by the 2^−ΔΔCt^ method with GAPDH or β-actin as the reference gene, reporting fold-change versus the indicated control condition

### Luciferase assay

C2C12 cells were co-transfected with the utrophin-A 5′UTR firefly luciferase reporter (5′Luc) and pRL-TK (Renilla control, Promega, Cat. No.: E2241) using Lipofectamine® 2000 (Invitrogen, Cat. No.: 11668019) in Opti-MEM® media (Gibco^TM^, Cat. No.: 31985062). After 24 hours, the media was refreshed, and luciferase activity was measured using the Dual-Glo® Luciferase Assay System (Promega, Cat. No.: E2920). Firefly luciferase signals were normalized to Renilla luciferase for transfection control (Ghosh and Basu 2015).

### RNA isolation from nuclear and cytosolic fractions

C2C12 cells were trypsinized at 37°C for 2 minutes and pelleted by centrifugation at 4000 rpm for 5 minutes. The cell pellets were washed three times with ice-cold 1XPBS, centrifuging at 4000 rpm for 5 minutes after each wash. The washed pellet was resuspended in RSB buffer (10 mM Tris-HCl pH 7.4, 10 mM NaCl, 3 mM MgCl_2_), incubated on ice for 5 minutes, and then centrifuged at 4000 rpm at 4°C for 5 minutes to obtain swollen cell pellets. The swollen pellets were gently resuspended in four volumes of RSBG40 buffer (10 mM Tris-HCl pH 7.8, 10 mM NaCl, 3 mM MgCl_2_, 10% glycerol, 0.5% NP-40, 0.5 mM DTT, and 10 U/mL murine RNase inhibitor-NEB-M0314S) and incubated on ice for 5 minutes. The mixture was centrifuged at 7000 rpm for 5 minutes at 4°C, and the supernatant was collected as the cytoplasmic fraction. The pellet was then resuspended in RSBG40 buffer, and to this suspension, 3.3% (w/w) sodium deoxycholate and 6.6% (v/v) Tween-40 (calculated as 1/10th the volume of the resuspension) were added. After 10 minutes of incubation on ice, the sample was centrifuged at 7000 rpm for 5 minutes at 4 °C. The resulting supernatant was collected as the nuclear fraction. To both cytoplasmic and nuclear fractions, 1 mL of TRIzol^TM^ reagent (Invitrogen, Cat. No.: 15596026) was added for RNA extraction, following the manufacturer’s protocol (Ghosh et al. 2021; Wang et al. 2006).

### Protein isolation from the nuclear and cytosolic fractions

C2C12 cells were harvested by trypsinization and pelleted by centrifugation at 500 × g for 5 minutes at 4 °C. The cell pellet was washed twice with ice-cold PBS and resuspended in ice-cold hypotonic lysis buffer (10 mM Tris-HCl, pH 7.9; 10 mM KCl; 1.5 mM MgCl_2_; 0.5 mM DTT; 0.1% NP-40; 1× protease inhibitor cocktail) at a ratio of 200 μL buffer per 10⁶ cells. The suspension was incubated on ice for 10 minutes to allow cell swelling. Cells were then lysed by gentle pipetting and centrifuged at 800 × g for 5 minutes at 4 °C. The supernatant, containing the cytoplasmic protein fraction, was collected and stored on ice. The nuclear pellet was washed once with hypotonic buffer and resuspended in nuclear extraction buffer (20 mM Tris-HCl, pH 7.9; 400 mM NaCl; 1 mM EDTA; 1 mM EGTA; 1 mM DTT; 1X protease inhibitor cocktail). The pellet was incubated on ice for 30 minutes with intermittent vortexing to promote nuclear lysis. Following incubation, the mixture was centrifuged at 13,000 rpm for 15 minutes at 4 °C. The supernatant, containing the nuclear protein fraction, was carefully collected and stored at −80 °C until use. Protein concentrations in both fractions were determined using the Bradford assay (Ghosh et al. 2021).

### Statistical analysis

All statistical comparisons were carried out using two-tailed Student’s t-tests, applied to pairwise contrasts. Tests were unpaired and used an alpha level of 0.05. For each experiment, the unit of replication (n) refers to independent biological experiments and technical replicates within an experiment were averaged to yield a single value per biological replicate before statistics. Data are presented as mean ± SEM. Normalized values were used where appropriate (e.g., Firefly/Renilla for luciferase assays; 2^−ΔΔCt^ for qPCR), and fold-change values are reported relative to the indicated control condition. Significance reporting follows the scheme used consistently across the experiments: “ns” denotes p > 0.05; * denotes p < 0.05; ** denotes p < 0.001; and *** denotes p < 0.0001.

## Acknowledgements

The authors thank the Council for Scientific and Industrial Research (CSIR) for providing fellowship to D.P. The authors also thank Anusandhan National Research Foundation (ANRF), Department of Science C Technology, Government of India (ANRF CRG/2022/007730) for providing financial support. The authors also thank the HPC facility and the cell culture facility of the S.N. Bose Innovation Centre of the University of Kalyani, DST-PURSE II, and the Personal Research Grant of the University of Kalyani. The authors thank Dr. Arunima Biswas, Assistant Professor, Department of Zoology, for providing the goat anti-mouse IgG-FITC antibody. The authors acknowledge the Central Instrumental Facility at the Department of Biological Sciences, IISER Kolkata, for providing the ITC facility.

## Data Availability Statement

All the data related to this article has been provided and will be available from the corresponding author upon request.

## Author Contributions

D.P.: Conceptualization, data curation, formal analysis, investigation, methodology, software, validation, visualization, and writing – original draft. A.B.: Methodology, software, validation, visualization, and writing – original draft. S.D., G.G., S.P., and S.S.: Formal analysis and investigation. U.B.: Conceptualization, funding acquisition, project administration, and writing – review and editing. All the authors finalized this version of the manuscript.

